# Fetal liver hepcidin secures iron stores in utero

**DOI:** 10.1101/799304

**Authors:** Lara Kämmerer, Goran Mohammad, Magda Wolna, Peter A. Robbins, Samira Lakhal-Littleton

## Abstract

In the adult, the liver-derived hormone hepcidin (HAMP) controls systemic iron levels by blocking the iron-exporting protein ferroportin (FPN) in the gut and spleen, the sites of iron absorption and recycling respectively. Impaired HAMP expression or FPN responsiveness to HAMP result in iron overload. HAMP is also expressed in the fetal liver but its role in controlling fetal iron stores is not understood. To address this question in a manner that safeguards against the confounding effects of altered maternal iron homeostasis, we generated fetuses harbouring a paternally-inherited ubiquitous knock-in of the HAMP-resistant *fpn*C326Y. Additionally, to safeguard against any confounding effects of altered placental iron homeostasis, we generated fetuses with a liver-specific knock-in of *fpn*C326Y or knockout of the *hamp* gene. These fetuses had reduced liver iron stores, and markedly increased FPN in the liver, but not in the placenta. Thus, in contrast to the adult, fetal liver HAMP operates cell-autonomously to increase fetal liver iron stores. Our findings also suggest that FPN in the placenta is permissive rather than regulatory of iron transport.

## Introduction

Iron is essential for growth and development. Suboptimal iron availability at birth is associated with cognitive, behavioural and motor skill deficits (1-5). These developmental deficits are not reversible by oral iron therapy in the neonate (6-13). In the first 6-9 months of age, the neonate is not fully competent in regulating dietary iron absorption in response to iron needs (14,15), and depends on liver iron stores to support its growth and development (16,17).

In the adult, the liver-derived hormone HAMP controls systemic iron availability, which in turn controls liver iron stores (18, 19). HAMP operates by blocking the iron export protein ferroportin (FPN) in the gut and the spleen, the sites of iron absorption and recycling respectively (18,19). Disruption of the HAMP/FPN axis, in a manner that reduces HAMP levels or FPN responsiveness to HAMP, leads to the iron-overload disease hemochromatosis. This is characterised by increased serum iron concentration, depletion of iron from the spleen and excess deposition in the liver (20). The hemochromatosis phenotype is recapitulated in adult mice harbouring ubiquitous knock-in of the HAMP-resistant *fpn*C326Y allele, or liver-specific knockout of the *hamp* gene (21, 22).

In contrast to adult HAMP, the role of fetal HAMP in determining liver iron stores in utero is not entirely understood. One study reported that endogenous HAMP was not expressed in the mouse fetal liver but that transgenic ubiquitous overexpression of the *hamp* gene resulted in anaemic fetuses (23). Another study found that ubiquitous homozygous loss of the *matriptase-2* gene in fetuses (matriptase-2 is a negative regulator of *hamp* expression) resulted in reduced body non-heme iron levels (24). A third study comparing fetuses harbouring a homozygous deletion of the hemochromatosis gene *hfe* (HFE is a protein that is necessary for *hamp* expression), with wild type littermates did not find any difference in total liver iron stores, but reported higher levels of non-heme iron in *hfe*-ko fetuses when mothers were fed an iron-rich diet. (25). Based on these studies, it has been hypothesised that fetal liver HAMP controls iron availability to the fetus by blocking FPN-mediated iron transport into the fetal capillaries of the placental syncytiotrophoblast (STB) (26).

A common feature of the above studies is that the genetic variants in the fetuses were also present in the fetal part of the placenta, in the mother, and in the maternal part of the placenta. We sought to interrogate the role of fetal liver HAMP without these potentially confounding effects. To avoid those associated with altered iron control in the mother or maternal side of the placenta, we generated fetuses harbouring a paternally-inherited ubiquitous knock-in of the HAMP-resistant *fpn*C326Y allele. To avoid the additional potentially confounding effects of altered iron-control in the fetal part of the placenta, we also generated fetuses harbouring a liver-specific knock-in of the HAMP-resistant *fpn*C326Y allele, or liver-specific knockout of the *hamp* gene, through paternal inheritance of the hepatocyte-specific Albumin-Cre recombinase transgene. For these models, we then compared the levels of liver iron stores between the affected fetuses and their littermate controls.

## METHODS

### Mice

All animal procedures were compliant with the UK Home Office Animals (Scientific Procedures) Act 1986 and approved by the University of Oxford Medical Sciences Division Ethical Review Committee. The conditional *fpn*C326Y^fl^ allele was generated as described previously (21). Females were mated between 9 and 12 weeks of age and fetuses were harvested from first pregnancies.

### Fetal tissue

Fetal tissues were harvested from PBS-perfused mothers at e13.5 or e17.5, washed further in PBS then either snap-frozen for RNA and iron measurement studies or fixed in formalin for histological studies.

### Immunostaining

Formalin-fixed paraffin-embedded (FFPE) tissue sections were stained with rabbit polyclonal anti-mouse FPN antibody (NBP1-21502, Novus biologicals) at 1/200 dilution. Alexa 488-conjugated anti-rabbit antibody (ab150073, Abcam) was then used as a secondary antibody at a dilution of 1/500.

### Iron indices

Determination of total elemental iron in tissues from PBS-perfused animals was carried out by inductively coupled plasma mass spectrometry (ICP-MS) as described previously (22). Haemoglobin was recorded from fresh blood using the HemoCue Hb 201+ system. DAB-enhanced Perls’ iron stain was carried out in FFPE sections as described previously (22).

### Diet provision during pregnancy

Unless otherwise stated, mothers were fed a standard chow diet containing 200ppm iron. In iron-loading studies, mothers were fed a diet containing 5000ppm iron (Teklad TD.140464) as soon as mated.

### Gene expression

Gene expression was measured by quantitative real-time PCR, using Applied Biosystems Taqman gene expression assay probes for fpn, tfr1, hamp and house-keeping gene β-Actin (Life Technologies, *Carlsbad, CA*). The CT value for the gene of interest was first normalised by deducting CT value for β-Actin to obtain a delta CT value. Delta CT values of test samples were further normalised to the average of the delta CT values for control samples to obtain delta delta CT values. Relative gene expression levels were then calculated as 2^-delta deltaCT^.

### Statistics

Values are shown as mean± standard error of the mean (S.E.M). Paired comparisons were performed using Student’s T test. R values reported are Pearson correlation coefficients.

## RESULTS AND DISCUSSION

### Fetal liver iron concentration and hepcidin expression increase in the third trimester

In the mouse, the fetal liver forms in the second half of gestation. We found that the total concentration of elemental iron in the fetal liver, measured by inductively coupled plasma mass spectrometry (ICP-MS) as described previously (21), increased by 2.75 fold between e13.5 and e17.5, to levels well above those in the maternal liver at e17.5 (and similar to those in non-pregnant females), consistent with increased liver iron stores (Figure 1A). Over the same period of time, fetal liver hamp mRNA expression increased by 24 fold (Figure 1B). Nevertheless, at e17.5, fetal liver hamp mRNA expression was ∼ 18 fold lower than that of maternal liver hamp (Figure 1B). As expected, and consistent with a role of iron stores in supporting growth, there was a positive correlation between the concentration of iron in the fetal liver and fetal weight at e17.5 (Figure 1C). Additionally, there was a positive correlation between the concentration of iron and the expression of hamp mRNA in the fetal liver at e.17.5 (Figure 1D).

**Figure 1:**
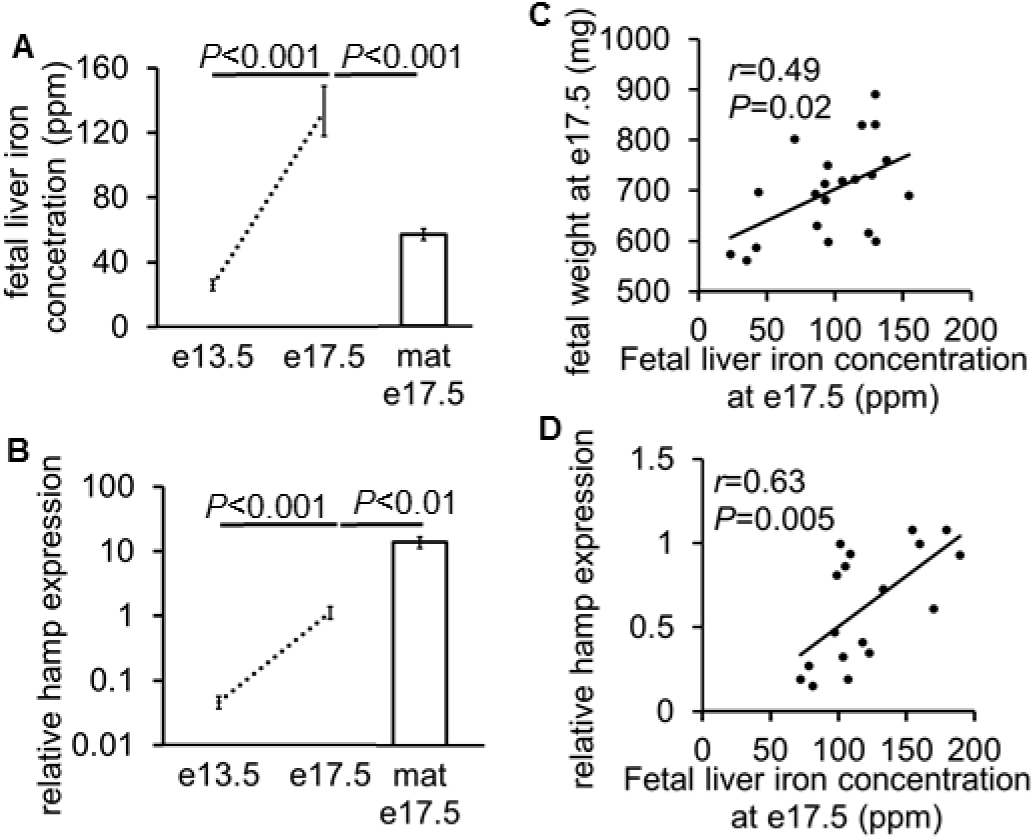
Fetal liver iron concentration and hepcidin expression increase in the third trimester. **A.** Total iron concentration in fetal livers harvested at e13.5 (n=27) and e17.5 (n=28) and in maternal livers at e17.5 (n=5). **B**. Relative hamp mRNA expression in fetal livers harvested at e13.5 (n=6) and e17.5 (n=6) and in maternal livers at e17.5 (n=6). **C.** Correlation between fetal weight and fetal liver iron concentration at e17.5 (n=21). **D**. Correlation between relative hamp expression and total iron concentration in fetal livers at e17.5 (n=18). Values are shown as mean±S.E.M. P values are calculated using Student’s t test, *r* is Pearson’s correlation coefficient.

### Ubiquitous loss of FPN responsiveness to hepcidin reduces fetal liver iron stores and haemoglobin at birth

In the adult, liver HAMP regulates liver iron stores (18-20). To probe the role of fetal HAMP in regulating fetal liver iron stores, we used mice harbouring a ubiquitous knock-in of a HAMP-resistant FPN. To guard against a confounding effect of maternal iron overload, we set up matings using wild type *fpn*^wt/wt^ mothers and *fpn*^wt/C326Y^ fathers heterozygous for the knock-in mutation of *fpn*C326Y, an isoform of FPN with intact iron export function but which is resistant to HAMP-mediated degradation. Adult mice harbouring this mutation develop the iron overload disease hemochromatosis, characterised by increased liver iron stores and depletion of iron from the spleen, by 12 weeks of age (21). In contrast to the adult mice, *fpn*^wt/C326Y^ fetuses at e17.5, had ∼30% lower fetal liver iron concentration compared with *fpn*^wt/wt^ littermate controls (Figure 2A). This was further confirmed histologically by DAB-enhanced Perls’ iron stain (Figure 2B). At birth (d0), liver iron concentration remained lower in *fpn*^wt/C326Y^ fetuses than in *fpn*^wt/wt^ littermate controls (Figure 2A). This difference disappeared within the first week of life, with mice of both genotypes having comparable liver iron stores at 1 and 4 weeks of age (Figure 2A). By 12 weeks of age, *fpn*^wt/C326Y^ mice had liver iron concentrations that were 115% higher than those of *fpn*^wt/wt^ littermate controls (Figure 2A), consistent with the hemochromatosis phenotype (21). Reduced liver iron stores in utero and at birth could not be attributed to re-distribution of iron into extra-hepatic tissues, as iron concentration in the remaining carcass (fetus-liver) of *fpn*^wt/C326Y^ fetuses was still lower than in *fpn*^wt/wt^ littermates at e17.5 (Figure 2C). Splenic iron concentration was comparable between mice of the two genotypes at birth (d0), and at 1 and 4 weeks of age, but lower in *fpn*^wt/C326Y^ mice than in *fpn*^wt/wt^ littermate controls at 12 weeks of age (Figure 2C), consistent with the hemochromatosis phenotype (21). Limited iron availability affects hemoglobin synthesis, and consistent with this, hemoglobin levels at birth (d0) were 16% lower in *fpn*^wt/C326Y^ mice than in *fpn*^wt/wt^ littermate controls (Figure 2D). By one week of age, hemoglobin levels were comparable between animals of the two genotypes (Figure 2D). Thus, ubiquitous loss of FPN responsiveness to hepcidin in fetal tissues results in reduced fetal liver iron stores and lower haemoglobin at birth.

**Figure 2:**
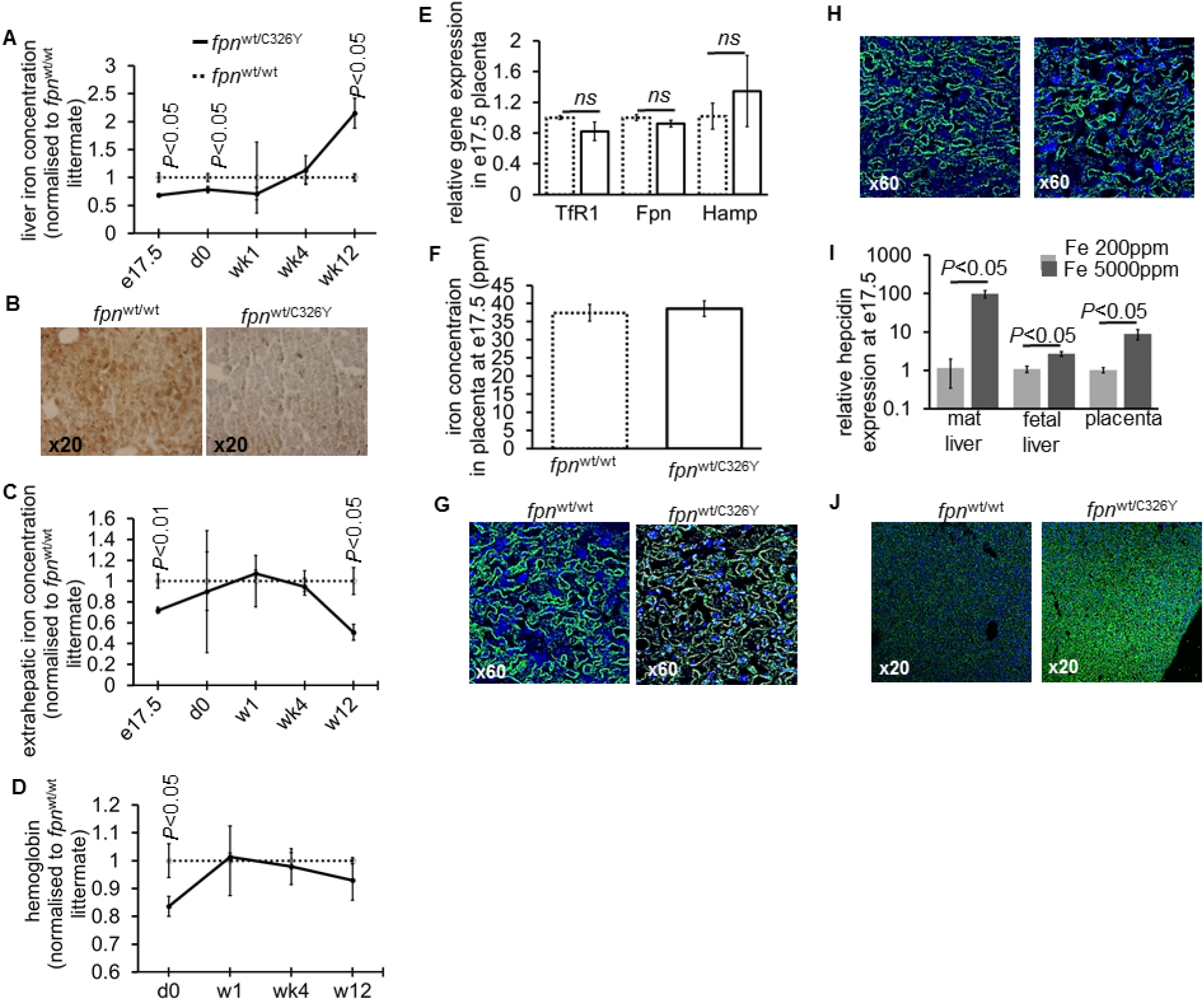
Ubiquitous loss of FPN responsiveness to hepcidin in the fetus reduces fetal liver iron stores and hemoglobin at birth. **A.** Liver iron concentration in *fpn*^wt/C326Y^ animals relative to *fpn*^wt/wt^ littermate controls at e17.5 (n=4, 8 respectively), day 0 (n=14,17), weekl (n=4, 6), week 4 (n=3, 3) and week 12 (n=3, 3) of age. *P* values are shown relative to *fpn*^wt/C326Y^ littermate controls at the respective timepoint. **B**. Representative images of DAB-enhanced Peris’ iron stain in the livers of *fpn*^wt/C326Y^ animals and *fpn*^wt/wt^ littermate controls at e17.5. **C**. Extra-hepatic iron concentration in *fpn*^wt/C326Y^ animals relative to *fpn*^,wt/wt^ littermate controls at e17.5 (carcass-liver) (n=6,11) and in the spleens of mice at day 0 (n=3,4), week 1 (n=3, 4), week 4 (n=3 per group) and week 12 (n=3 per group) of age. P values are shown relative to *fpn*^wt/wt^ littermate controls at the respective timepoint. **D**. Hemoglobin concentration in *fpn*^wt/C326Y^ animals relative to *fpn*^wt/wt^ littermate controls at day 0 (n=6, 11), weekl (n=3, 4), week 4 (n=3 per group) and week 12 (n=3 per group) of age. **E.** Relative expression ofTfRI, Fpn and Hamp mRNA transcripts in placentae of *fpn*^wt/C326Y^ fetuses (n=5) and *fpn*^wt/wt^ littermates (n=3) at e17.5. **F**. Iron concentration in placentae of/*fpn*^wt/C326Y^ fetuses (n=11) and *fpn*^wt,wt^ littermates (n=6) at e17.5. **G**. Representative images of FPN immunostaining in the STB from placentae of *fpn*^wt/C326Y^ animals and *fpn*^wt/wt^ littermate controls at e17.5. **H**. Representative images of FPN immunostaining in the STB of placentae harvested at e17.5 from *fpn*^wt/wt^ mothers carrying *fpn*^wt/wt^ litters and fed a normal iron diet (200ppm) or an iron-rich diet (5000ppm). **I**. Relative expression of Hamp mRNA transcript in maternal livers (n=3 per group), fetal livers (n=6 per group) and corresponding placentae (n=6 per group) harvested at e17.5 from mothers fed a normal iron diet (200ppm) or an iron-rich diet (5000ppm). **J**. Representative images of FPN immunostaining in the livers of *fpn*^wt/C326Y^ animals and *fpn*^wt/wt^ littermate controls at e17.5. Values are shown as mean±S.E.M. *P* values are calculated using Student’s t test.

Next, we sought to determine whether the reduction in fetal iron stores in *fpn*^wt/C326Y^ animals could be attributed to changes in placental iron handling. However, we found that placental iron concentration, and the expression of iron-regulated transcripts *Fpn, transferrin receptor 1* (TfR1) and *hamp* were all comparable between placentae of *fpn*^wt/C326Y^ mice and placentae of *fpn*^wt/wt^ littermates (Figure 2E-F), suggesting that placental iron handling is not different between animals of the two genotypes. When we examined the STB, we found that FPN levels were not visibly different between placentae of *fpn*^wt/C326Y^ mice and placentae of *fpn*^wt/wt^ littermates (Figure 2G), indicating FPN at this site is not actively regulated by hepcidin. Consistent with this, we found that provision of an iron-loaded diet to *fpn*^wt/wt^ mothers carrying *fpn*^wt/wt^ litters failed to alter FPN in the placental STB (Figure 2H), despite concomitantly increasing the expression of hepcidin in the fetal and maternal livers and in the placenta itself (Figure 2I).Thus altered placental iron handling does not explain reduced liver iron stores seen in *fpn*^wt/C326Y^ fetuses. Apart from the placenta, the other site of FPN expression is the fetal liver itself. Therefore, we examined whether FPN expression in the livers of *fpn*^wt/C326Y^ fetuses was altered, and found that it was markedly increased compared to livers of *fpn*^wt/wt^ littermates at e17.5 (Figure 2J). Therefore, we hypothesised that this marked increase in FPN expression is the cause of decreased fetal liver iron stores seen in *fpn*^wt/C326Y^ fetuses.

### Liver-specific loss of FPN responsiveness to hepcidin reduces fetal liver iron stores

To test the hypothesis that increased FPN in the fetal liver leads to a reduction in iron stores, we generated fetuses harbouring a liver-specific knock-in of the *fpn*C326Y allele, by mating *fpn*C326Y^fl/fl^ mothers with *fpn*C326Y^fl/fl^ fathers transgenic for Cre recombinase driven by the hepatocyte-specific albumin promoter (*fpn*C326Y^fl/fl^, Alb.Cre+). We then harvested *fpn*C326Y^fl/fl^, Alb.Cre+ fetuses and *fpn*C326Y^fl/fl^ littermates at e17.5, and found that *fpn*C326Y^fl/fl^, Alb.Cre+ fetuses had a 26% reduction in liver iron concentration relative to *fpn*C326Y^fl/fl^ littermates (Figure 3A).This was confirmed histologically by DAB-enhanced Perls’ iron stain (Figure 3B). Additionally, *fpn*C326Y^fl/fl^, Alb.Cre+ fetuses had increased hepatocyte FPN expression compared with *fpn*C326Y^fl/fl^ littermates at e17.5 (Figure 3C). These results confirm that FPN in fetal hepatocytes is subject to regulation by HAMP and that this regulation is important for the control of fetal liver iron stores.

**Figure 3:**
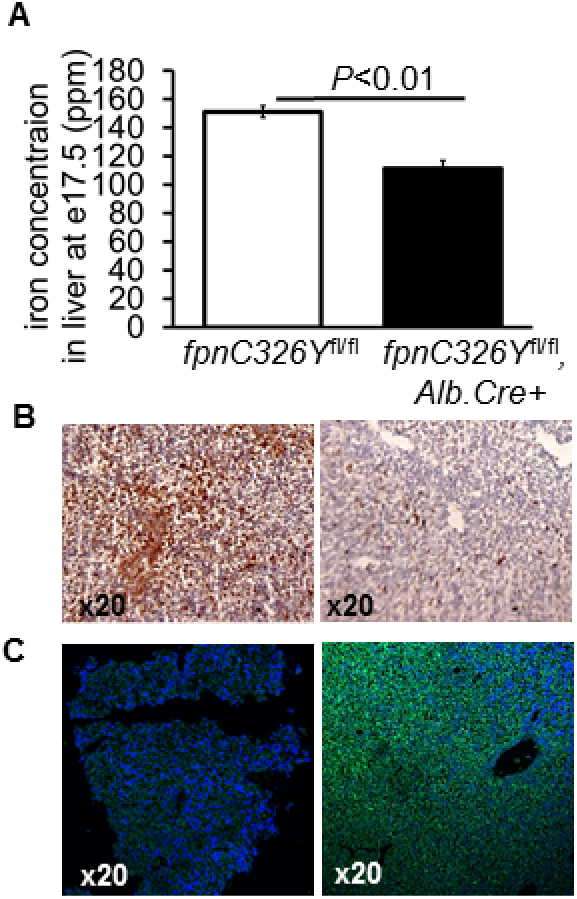
Liver-specific loss of FPN responsiveness to hepcidin reduces fetal liver iron stores. **A.** Total Iron concentration in the livers of *fpnC326Y*^fl\fl^ Alb.Cre+ animals and *fpnC326 Y*^fl\fl^ littermate controls at e17.5 (n=3 per group). **B**. Representative images of DAB-enhanced Peris’ iron stain in the livers of *fpnC326 Y*^fl\fl^, Alb.Cre+ animals and *fpnC326 Y*^fl\fl^ littermate controls at e17.5. **C**. Representative images of FPN immunostaining in the livers of *fpnC326 Y*^fl\fl^, Alb.Cre+ animals and *fpnC326 Y*^fl\fl^ littermate controls at e17.5. Values are shown as mean±S.E.M. *P* values are calculated using Student’s t test.

### Fetal liver hepcidin regulates fetal liver FPN and iron stores

Next, we set out to identify the source of HAMP that regulates fetal liver FPN and consequently its iron stores. The previous observations that fetal liver hamp mRNA increased 24 fold between e13.5 and e17.5 and that it correlated positively with fetal liver iron stores (Figures 1B, D) suggest that the fetal liver is the source of HAMP that regulates fetal liver FPN and iron concentration. To test this hypothesis, we deleted the *hamp* gene specifically in the fetal liver by mating *hamp*^fl/fl^ mothers with *hamp*^fl/fl^, Alb.Cre+ fathers. We found that at e17.5, hamp mRNA expression was reduced by 80% in the livers of *hamp*^fl/fl^, Alb.Cre+ fetuses relative to *hamp*^fl/fl^ littermates, confirming the activity of the Cre allele at this gestational stage (Figure 4A). Loss of fetal liver HAMP resulted in a 47% decrease in fetal liver iron concentration compared with *hamp*^fl/fl^ littermates (Figure 4B), and this was further confirmed histologically by DAB-enhanced Perls’ iron stain (Figure 4C). In contrast, adult *hamp*^fl/fl^, Alb.Cre+ mice had 19 fold increase in liver iron concentration compared with *hamp*^fl/fl^ controls, consistent with the well-recognised role for hepatic HAMP in controlling liver iron stores in the adult (Figure 4D). When we compared FPN expression in *hamp*^fl/fl^ and *hamp*^fl/fl^, Alb.Cre+ fetuses at e17.5, we found that FPN expression was markedly increased in the livers of *hamp*^fl/fl^, Alb.Cre+ fetuses (Figure 4E), but comparable between placentae of the two genotypes (Figure 4F).

**Figure 4:**
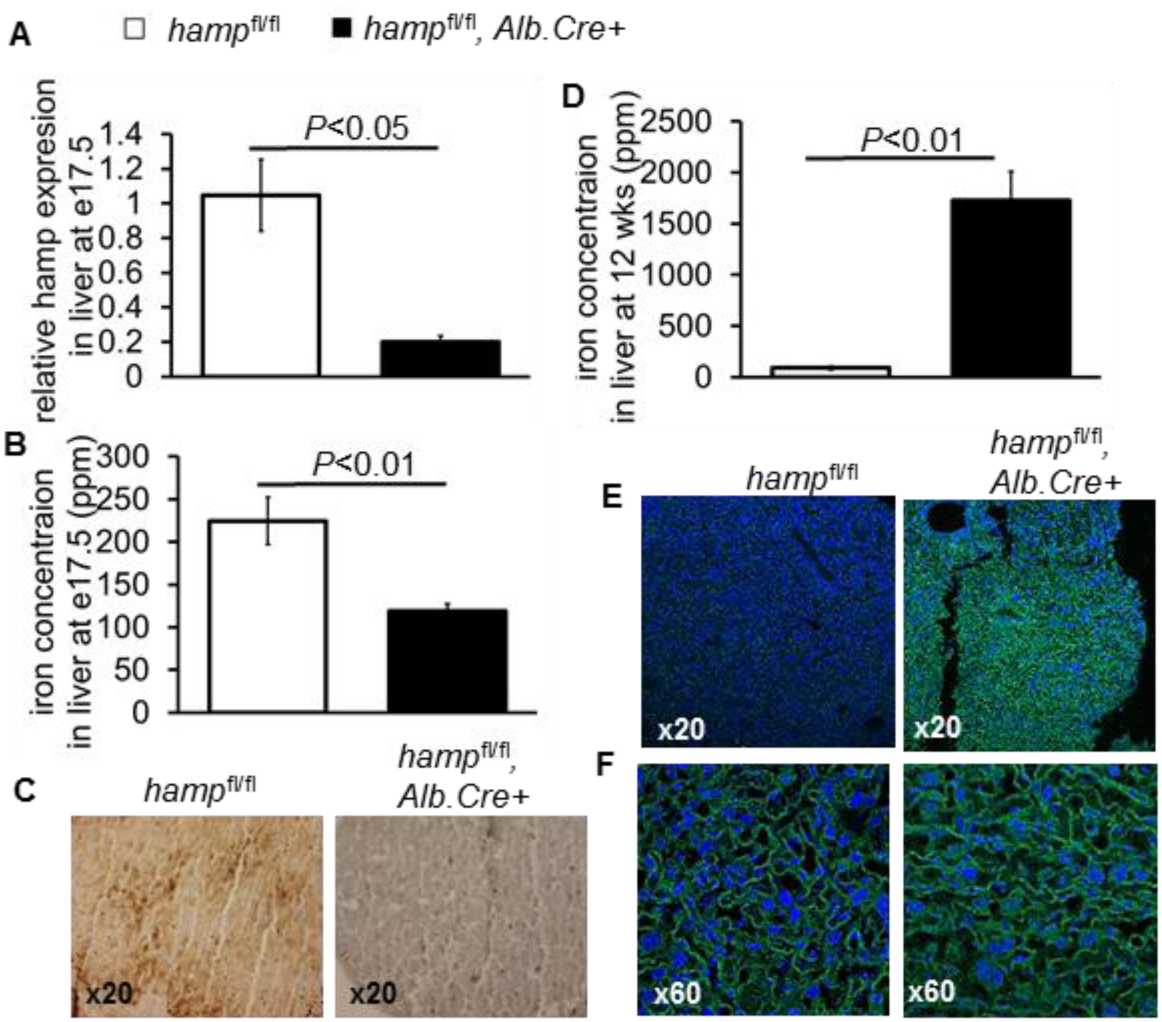
Liver-specific loss of hepcidin reduces fetal liver iron stores. **A**. Relative hamp mRNA expression in livers of *hamp*^fl\fl^, Alb.Cre+ animals (n=5) and *hamp*^fl\fl^ littermate controls (n=4) at e17.5. **B**. Total iron concentration in the livers of *hamp*^fl\fl^, Alb.Cre+ animals and hamp^m^ littermate controls at e17.5 (n=8 per group). **C**. Representative images of DAB-enhanced Peris’ iron stain in the livers of *hamp*^fl\fl^, Alb.Cre+ animals and *hamp*^fl\fl^ littermate controls at e17.5. **D**. Total iron concentration in the livers of *hamp*^fl\fl^, Alb.Cre+ animals and *hamp*^fl\fl^ littermate controls at 12 weeks of age (n=3 per group). **E**. Representative images of FPN immunostaining in the livers of *hamp*^fl\fl^, Alb.Cre+ animals and *hamp*^fl\fl^ littermate controls at e17.5. **F**. Representative images of FPN immunostaining in the STB from placentae of *hamp*^fl\fl^, Alb.Cre+ animals and *hamp*^fl\fl^ littermate controls at e17.5.Values are shown as mean±S.E.M. *P* values are calculated using Student’s t test.

The widely accepted consensus is that fetal liver HAMP regulates iron availability to the fetus by inhibiting placental FPN (23). This consensus is based on animal studies in which the iron homeostatic machinery is altered in the fetus, the mother and in the fetal and maternal parts of the placenta (24-26). By using paternally inherited alleles, our studies eliminate the confounding effects of altered iron homeostasis in the mother and in the maternal side of the placenta. The use of paternally-inherited liver-specific Albumin-Cre recombinase to disrupt the HAMP/FPN axis in the fetal liver further eliminates any confounding effects of altered iron homeostasis in the fetal part of the placenta. The important finding from these studies is that fetal liver HAMP operates in a cell-autonomous manner to secure fetal liver iron stores. The finding that fetal liver HAMP expression is significantly lower than that of maternal liver HAMP is consistent with an autocrine rather than endocrine mode of action. Liver iron endowment at birth is important for supporting neonatal growth, and indeed these stores are rapidly depleted in early life (16, 17). To ensure adequate liver iron endowment at birth, there is a rapid build-up of iron levels within the fetal hepatocytes. Normally, this would be prevented by the action of Iron Regulatory Proteins IRPs, that act to stabilise the *fpn* transcript, thus increasing FPN and cellular iron efflux (27). The cell autonomous blockade of FPN protein by fetal HAMP may well serve to counteract the action of IRPs in order to allow the rapid build-up of iron levels in hepatocytes.

Another important finding from our studies is that FPN in the STB is not subject to regulation by HAMP, suggesting that its role in this setting is permissive rather than regulatory. Further studies are warranted to determine the sites of FPN and HAMP expression in maternal and fetal cell populations within the placenta, and to understand the regulatory interactions between them. Such studies necessitate genetic tools that can target specific cell populations within the placenta without affecting iron control in the fetus or the mother.

## ACKNOWLEDGMENTS

This work was supported by a Medical Research Council project Grant (MR/L010054/1) awarded to S L-L and PAR. S L-L was the recipient of a British Heart Foundation Intermediate Basic Science Postdoctoral Fellowship (FS/12/63/29895).

## Notes

#### Summary of Updates

The manuscript abstract has been shortened to 180 words. The text in the body of the manuscript has been checked and altered gramatically in places. Additional citations were included.

